# Reciprocal connectivity of the periaqueductal gray with the ponto-medullary respiratory network in rat

**DOI:** 10.1101/2020.09.15.298927

**Authors:** Pedro Trevizan-Baú, Werner I. Furuya, Stuart B. Mazzone, Davor Stanić, Rishi R. Dhingra, Mathias Dutschmann

## Abstract

Synaptic activities of the periaqueductal gray (PAG) can modulate or appropriate the respiratory motor activities in the context of behavior and emotion via descending projections to nucleus retroambiguus. However, alternative anatomical pathways for the mediation of PAG-evoked respiratory modulation via core nuclei of the brainstem respiratory network remains only partially described. We injected the retrograde tracer Cholera toxin subunit B (CT-B) in the pontine Kölliker-Fuse nucleus (KFn, *n*=5), medullary Bötzinger (BötC, *n*=3) and pre-Bötzinger complexes (pre-BötC; *n*=3), and the caudal raphé nuclei (*n*=3), and quantified the ascending and descending connectivity of the PAG. CT-B injections in the KFn, pre-BötC, and caudal raphé, but not in the BötC, resulted in CT-B-labeled neurons that were predominantly located in the lateral and ventrolateral PAG columns. In turn, CT-B injections into the lateral and ventrolateral PAG columns (*n*=4) yield the highest numbers of CT-B-labeled neurons in the KFn and far fewer numbers of labeled neurons in the pre-BötC and caudal raphé. Analysis of the relative projection strength revealed that the KFn shares the densest reciprocal connectivity with the PAG (ventrolateral and lateral columns, in particular). Overall, our data imply that the PAG may engage a distributed respiratory rhythm and pattern generating network beyond the nucleus retroambiguus to mediate downstream modulation of breathing. However, the reciprocal connectivity of the KFn and PAG suggests specific roles for synaptic interaction between these two nuclei that are most likely related to the regulation of upper airway patency during vocalization or other volitional orofacial behaviors.

**Highlights:**

- The lateral and ventrolateral PAG project to the primary respiratory network.
- The Kölliker-Fuse nucleus shares the densest reciprocal connectivity with the PAG.
- The Bötzinger complex appears to have very little connectivity with the PAG.

## 1. Introduction

Stimulation of the lateral and ventrolateral cell columns of the midbrain periaqueductal gray (lPAG and vlPAG, respectively) can elicit behavioral modulations and appropriations of ongoing respiratory activity, in particular during vocalization (Magoun et al., 1937; Jürgens and Pratt, 1979, Larson and Kistler, 1984; Jürgens, 1994; Jürgens, 2009). In addition, these cell columns of the PAG have function in the mediation of cardio-respiratory effects associated with emotions (Carrive et al., 1997; Fanselow, 1991; Walker and Carrive, 2003). Thus, the PAG is seen as the primary interface that connects cortical and limbic forebrain circuits with autonomic networks in the brainstem (Dampney et al., 2013; Subramanian and Holstege, 2014).

The neuroanatomical framework for PAG-evoked modulation of cardio-respiratory activity is linked to PAG descending projections to the nucleus retroambiguus (NRA) in the caudal medulla oblongata (Holstege et al., 1989; Holstege et al., 1997; Vanderhorst et al., 2000). In turn, the NRA projects to motoneurons of the nucleus ambiguus (Vanderhorst et al., 2001) and to various pools of respiratory motoneurons in the ventral horn of the spinal cord (Vanderhorst and Holstege, 1995; Holstege et al., 1997). Therefore, the PAG-NRA pathway is considered essential for PAG-mediated volitional and emotional modulation of breathing (Subramanian et al., 2008a, b; Subramanian and Holstege, 2014; Faul et al., 2019). However, while the NRA has a role in respiratory rhythm generation (Jones et al., 2012), it does not contribute to the generation of the respiratory motor pattern (Jones et al., 2016a), which depends on synaptic interactions between the rhythmogenic pre-Bötzinger complex (pre-BötC; Smith et al., 1991; Feldman and Del Negro, 2006) in the rostral medulla, and pontine circuits (Jones et al., 2016b). Given that the PAG is not an integral part of the respiratory pattern generating circuit (Farmer et al., 2014), the mediation of respiratory modulation via the PAG-NRA pathway may not bypass the ponto-medullary rhythm and pattern generating network. Indeed, various anterograde and retrograde tracing studies have reported that descending PAG projections also target broader pontine and medullary brainstem areas, including the pontine tegmentum (Holstege, 1991), the rostral ventrolateral medulla (Carrive et al., 1988; Yasui et al., 1990), subretrofacial nucleus (Carrive et al.,1989; Carrive and Bandler, 1991), and the caudal raphé nuclei (Beitz et al., 1983; van Bockstaele et al., 1991; Cowie and Holstege, 1992). However, many of these studies were performed in cat and were often focused on general PAG pathways linked to cardiorespiratory regulation during defence behavior. The specific connectivity with core nuclei of the brainstem respiratory rhythm and pattern generating network were often not specifically investigated, except in the work of Krout et al. (1998) who reported a modest amount of anterograde-labeled fibres in the Kölliker-Fuse nucleus (KFn), a core nucleus of the pontine respiratory group (Dutschmann and Dick, 2012).

To develop a better understanding of the anatomical frameworks for the PAG-mediated modulation of breathing in the context of emotion and behavior, we specifically assessed the relative strength of the reciprocal connectivity of the lPAG and vlPAG with core nuclei of the respiratory network such as the Bötzinger complex (BötC), pre-BötC, KFn, and the caudal raphé nuclei using a retrograde tracing approach.

## 2. Results

We microinjected the retrograde tracer CT-B (150 nL) unilaterally in the pontine KFn (*n*=5), the medullary BötC(*n*=3) and pre-BötC(*n*=3), the caudal raphé (*n*=3), or the midbrain PAG (*n*=4). CT-B injections were anatomically confined to discrete injection sites in the brainstem target areas (see representative photographs, Fig. 1 & 3A). Note that the same tracer injections were reported in a most recent study from our laboratory that performed an extensive analysis of descending forebrain inputs to these brainstem nuclei (Trevizan-Baú et al., 2020).

**Figure 1.**
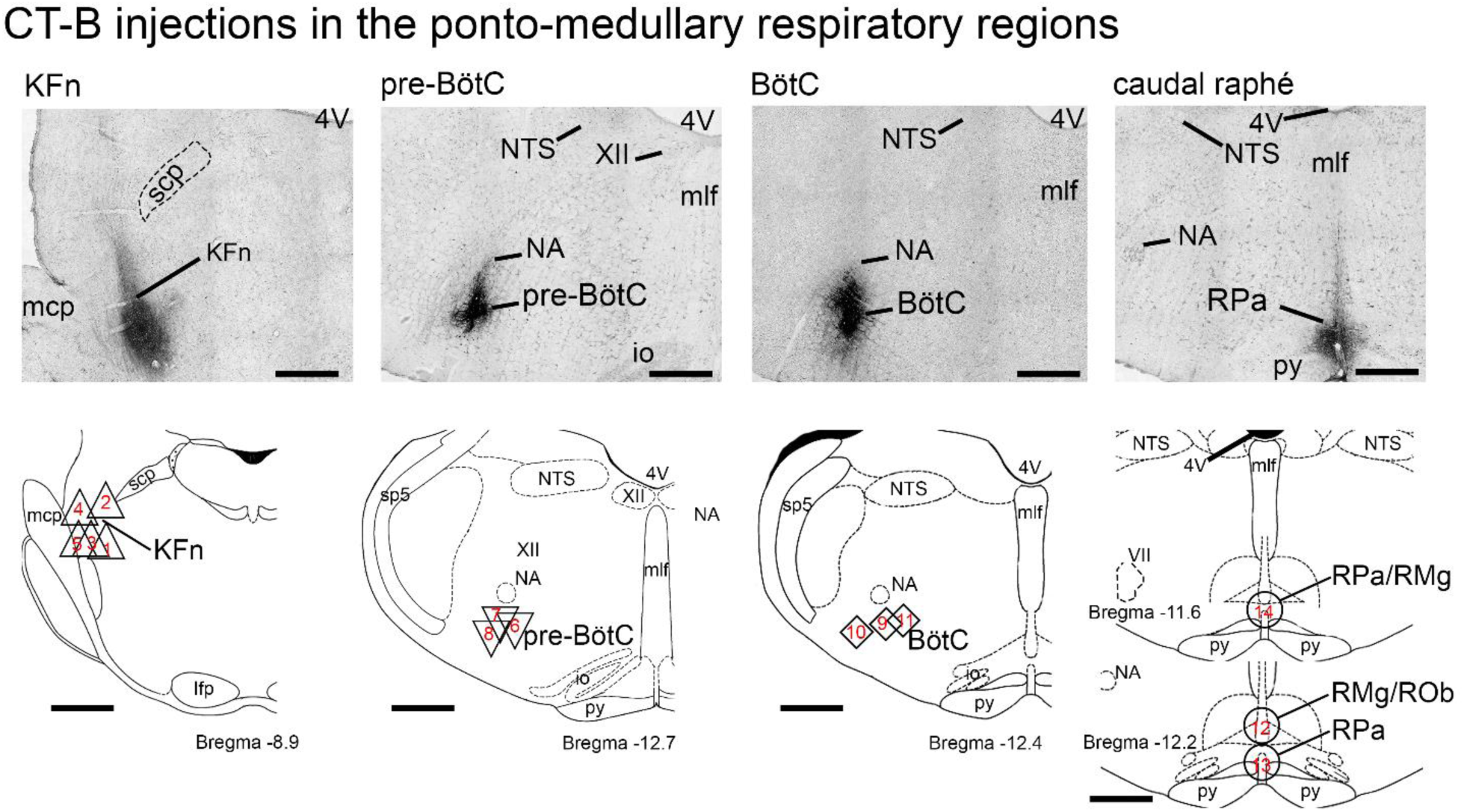
CT-B injections in the pontine Kölliker-Fuse nucleus, the medullary pre-Bötzinger complex and Bötzinger complex, and the caudal raphé. Photomicrographs of representative injections (top) and schematic drawings (bottom) depicting the location of all CT-B injections in the KFn (regular triangle), pre-BötC (upside-down triangle), BötC (square), caudal raphé (circles) relative to bregma. Each number represents a single experimental case. *Abbreviations*: 4V = fourth ventricle; BötC = Bötzinger complex; KFn = Kölliker-Fuse nucleus; mcp = middle cerebellar peduncle; io = inferior olive; mlf = medial longitudinal fasciculus; NA = nucleus ambiguous; NTS = solitary tract nucleus; PAG = periaqueductal gray; pre-BötC =, pre-Bötzinger complex; py = pyramidal tract; RMg = raphé magnus; Rob = raphé obscurus; RPa = raphé pallidus; scp = superior cerebellar peduncle; XII = facial nucleus. *Scale bars*: 200µm.

### 2.1. Distribution of retrogradely labeled neurons in the PAG after CT-B injections in ponto-medullary respiratory nuclei

The numbers of CT-B-retrogradely labeled cell bodies in the ipsilateral and contralateral midbrain PAG was 448±121 labeled neurons per case (n/c) after CT-B injection in the pontine KFn, and 303±98 n/c and 353±109 n/c after CT-B injection in the medullary pre-BötC and caudal raphé, respectively (Fig. 2A). However, no CT-B-labeled neurons were detected following tracer injection in the BötC (Fig. 2A). Analysis of the ipsilateral and contralateral distribution of CT-B-labeled neurons (Fig. 2A) revealed a significant ipsilateral dominance of descending PAG projections targeting the KFn (394±109 vs. 55±15 neurons/case (n/c); *p*<0.05), and a tendency of ipsilateral PAG projections to the pre-BötC (247±84 vs. 56±16 n/c). Because injections in the caudal raphé were in the midline of the medulla, we observed similar numbers of CT-B labeled neurons in the left and right hemisphere of the PAG (data not shown).

**Figure 2.**
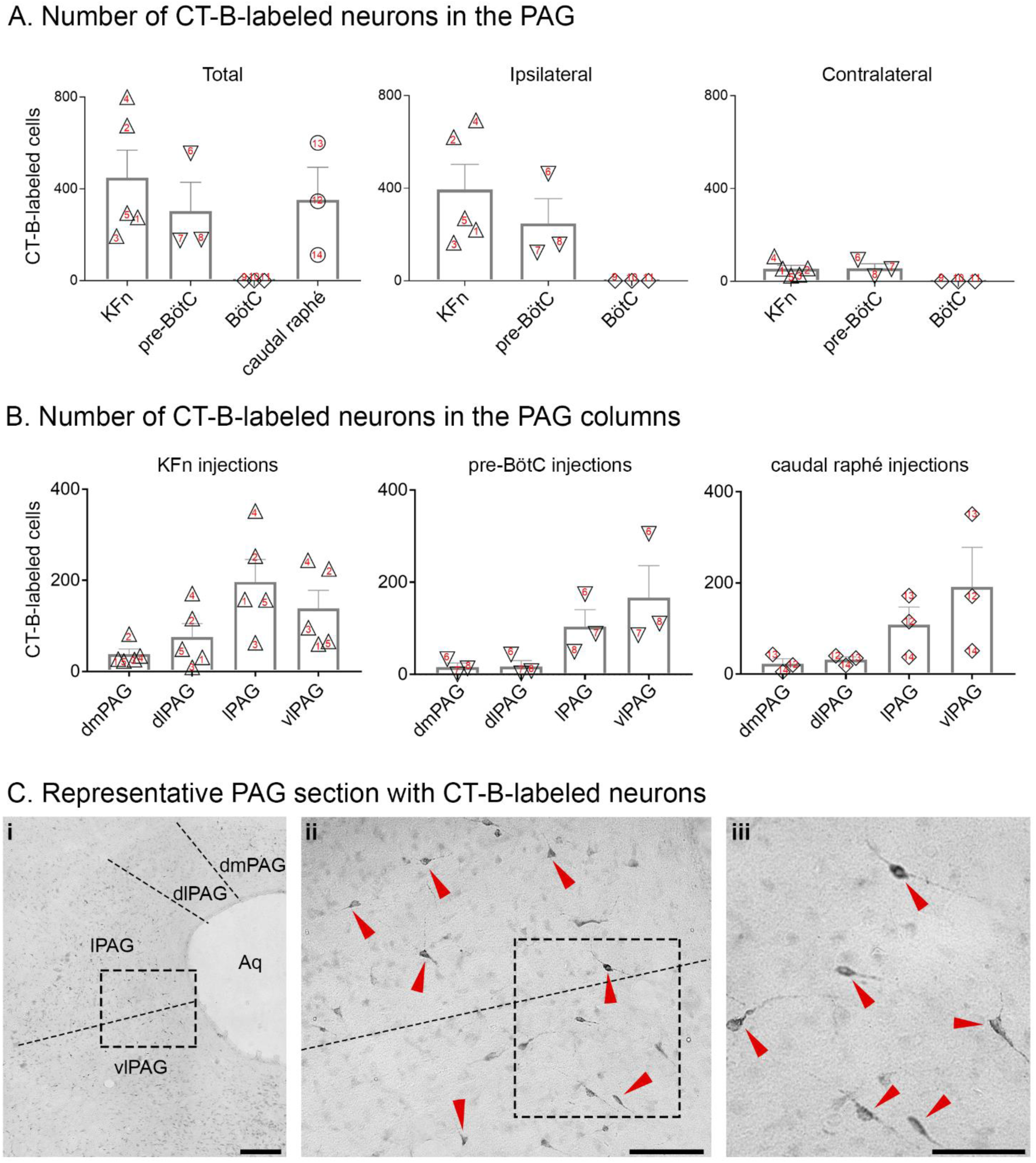
Injections in the ponto-medullary target regions resulted in CT-B-labeled neurons in the midbrain PAG. **A:** Total, ipsilateral, and contralateral distribution of labeled neurons in the PAG after injections in the KFn, pre-BötC, BötC, or caudal raphé. Analysis of the ipsilateral and contralateral distribution of CT-B-labeled neurons revealed a significant ipsilateral dominance of descending PAG projections targeting the KFn (*p*<0.05), and a tendency of ipsilateral PAG projections to the pre-BötC. **B:** Total number of CT-B-labeled neurons in the dmPAG, dlPAG, lPAG and vlPAG after CT-B injections in the pontine KFn, pre-BötC, or caudal raphé. Overall, we observed a predominance of CT-B-labeled neurons in the lateral and ventrolateral columns of the PAG. All values are expressed as mean ± standard error of the mean. **C:** Representative images of CT-B-labeled neurons in the PAG: i) the photomicrograph at the lowest magnification shows the retrograde labeled neurons in the context of the PAG columns; ii) at higher magnification (20x), CT-B-labeled neurons (red arrowheads) in the lPAG and vlPAG are shown; and iii) at 40x, CT-B-labeled neurons (red arrowheads) is more evident. *Abbreviations*: Aq = aqueduct; BötC = Bötzinger complex; KFn = Kölliker-Fuse nucleus; dlPAG = dorsolateral PAG; dmPAG = dorsomedial PAG; lPAG = lateral PAG; PAG = periaqueductal gray; vlPAG = ventrolateral PAG; pre-BötC = pre-Bötzinger complex. *Scale bars*: 100µm (i); 50µm (ii); 25µm (iii).

Analysis of the distribution of CT-B-labeled neurons in specific longitudinal cell columns of the PAG, which include the dorsomedial (dmPAG), dorsolateral (dlPAG), lateral (lPAG), and ventrolateral (vlPAG) columns (Bandler et al., 1991; Carrive, 1993), revealed that the majority of the projection neurons were located in the lPAG and vlPAG (Fig. 2B). Although CT-B-labeled neurons were observed throughout the rostrocaudal extension of the PAG columns (i.e., rostral, intermediate, and caudal PAG), the majority of the projection neurons for all experimental cases were located in the intermediate and caudal aspects of the respective PAG columns (data not shown).

### 2.2. Distribution of retrogradely labeled neurons in ponto-medullary respiratory nuclei after CT-B injections in the lateral and ventrolateral columns of the PAG

The locations of CT-B injections in the lPAG and vlPAG are illustrated in Fig 3A. Overall, our analysis revealed the highest number of CT-B-labeled neurons in the KFn, in comparison to the medullary BötC, pre-BötC, or caudal raphé (*p*<0.05; Fig. 3B, C). Specifically, injections in the PAG revealed on average 48±6 n/c in the KFn, versus only 6±1 n/c in the pre-BötC, 5±2 n/c in the BötC, or 1±1 n/c in the caudal raphé. Analysis of the laterality of retrogradely neurons in the pontine KFn indicated the highest number of CT-B-labeled neurons located ipsilaterally (*p*<0.05; Fig. 3B). We detected 32±3 neurons/case (n/c) in the ipsilateral KFn and 15±4 n/c in the contralaterally. In the medullary pre-BötC, we observed a tendency of higher number of CT-B-labeled neurons in the ipsilateral hemisphere. We detected 5±1 n/c located ipsilaterally and 1±1 n/c contralaterally (Fig. 3B). Finally, in the BötC, we detected 2±1 n/c in the ipsilateral hemisphere, and 3±1 n/c in the contralaterally (not significant; Fig. 3B). Please note that relative proportion of efferent versus afferent projections is an order of magnitude different because the entirety of a PAG column was not covered by a single CT-B injection, whereas all cells within a PAG column were counted when quantifying efferent projections. Given that the injection volumes were of a consistent volume, and were targeted toward the caudal aspect of the PAG in the case of afferent projections (see Fig 3A), their relative proportions should still be a reliable measure for comparison.

**Figure 3.**
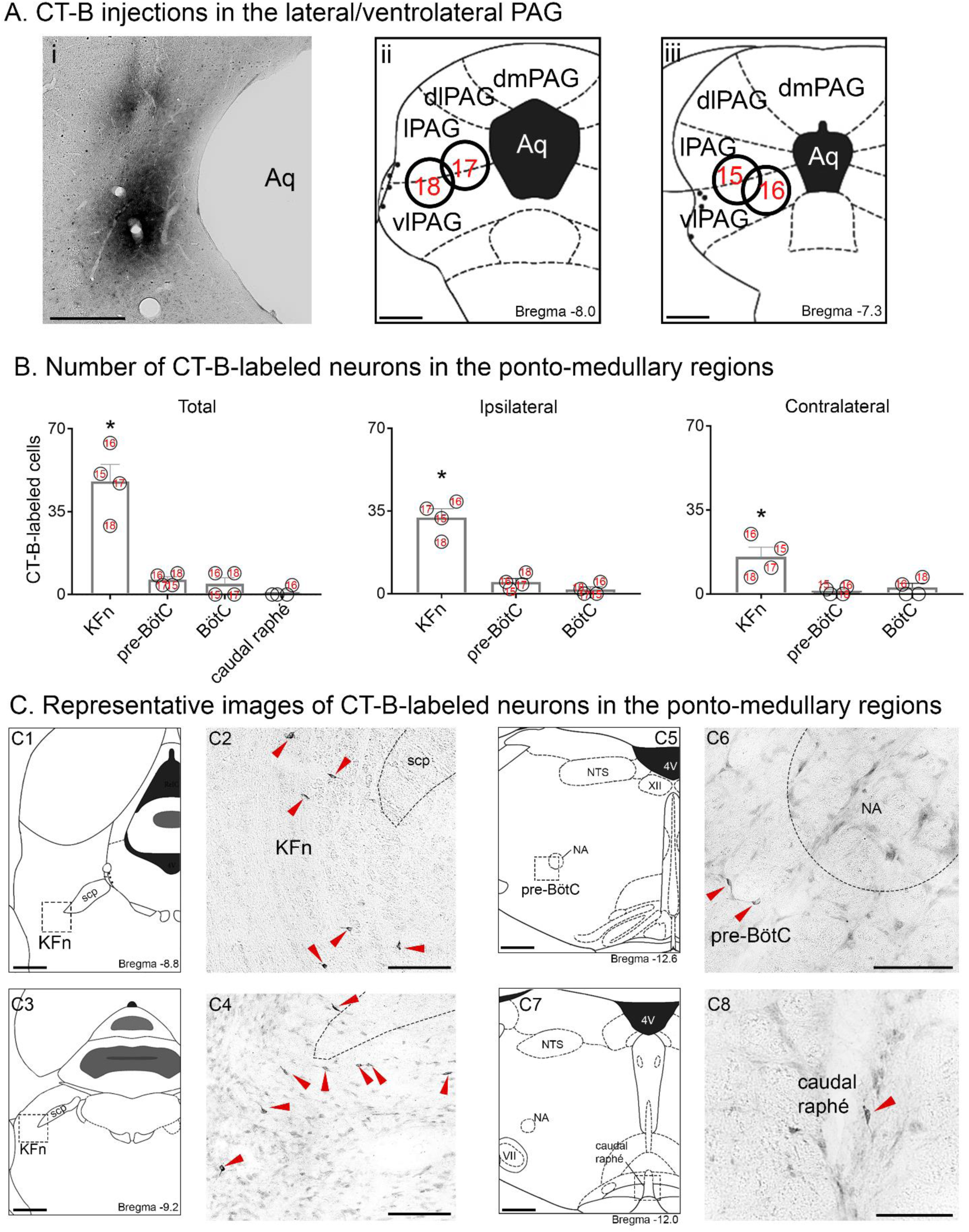
CT-B injections in the lateral/ventrolateral columns of the midbrain PAG resulted in CT-B-labeled neurons in the pontine KFn, the medullary pre-BötC and BötC, and the caudal raphé. **A:** Photomicrograph of a representative CT-B injection (i) and schematic drawings (ii, iii) depicting the location of all CT-B injections in the PAG (circles). Each number represents a single experimental case. **B:** Total, ipsilateral, and contralateral cell counts of CT-B-labeled neurons in the KFn, pre-BötC, BötC, and the caudal raphé after injections in the PAG. We observed a greater number of CT-B-labeled neurons in the KFn, compared to pre-BötC, BötC, and caudal raphé. Additionally, our analysis revealed the highest number of CT-B-labeled neurons in the ipsilateral KFn (*p*<0.05). In the medullary pre-BötC, we observed a tendency of higher number of CT-B-labeled neurons ipsilaterally, and no laterality was detected for the BötC. All values are expressed as mean ± standard error of the mean. **C:** Representative images of CT-B labeled neurons in the KFn (C1-4), pre-BötC (C5-6), and caudal raphé (C7-8). The photomicrograph on the right (C2, C4, C6, & C8) are of boxed regions marked in C1, C3, C5 and C7, respectively, to show the CT-B-labeled cells. *Abbreviations*: 4V = fourth ventricle; Aq = aqueduct; BötC = Bötzinger complex; dlPAG = dorsolateral PAG; dmPAG = dorsomedial PAG; io = inferior olive; KFn = Kölliker-Fuse nucleus; lPAG = lateral PAG; mcp = middle cerebellar peduncle; mlf = medial longitudinal fasciculus; NA = nucleus ambiguous; NTS = solitary tract nucleus; PAG = periaqueductal gray; pre-BötC = pre-Bötzinger complex; py = pyramidal tract; scp = superior cerebellar peduncle; vlPAG = ventrolateral PAG; XII = facial nucleus. *Scale bars*: 400µm (A; C1, C3, C5, & C7); 100µm (C2, C4, C6, & C8).

### 2.3. Summary of the afferent and efferent connectivity of lateral and ventrolateral PAG columns with respiratory nuclei in the ponto-medullary brainstem

The topographical organization of the descending and ascending PAG projections is summarized in a network connectivity graph (Figure 4). The graph shows the relative proportion of efferent projections from the midbrain PAG columns to each ponto-medullary target nuclei (Fig. 4A), and the afferent inputs to the PAG arising from the downstream targets (Fig. 4B). The functional implications of the reciprocal connectivity of the PAG in the context of volitional control of breathing are discussed below.

**Figure 4.**
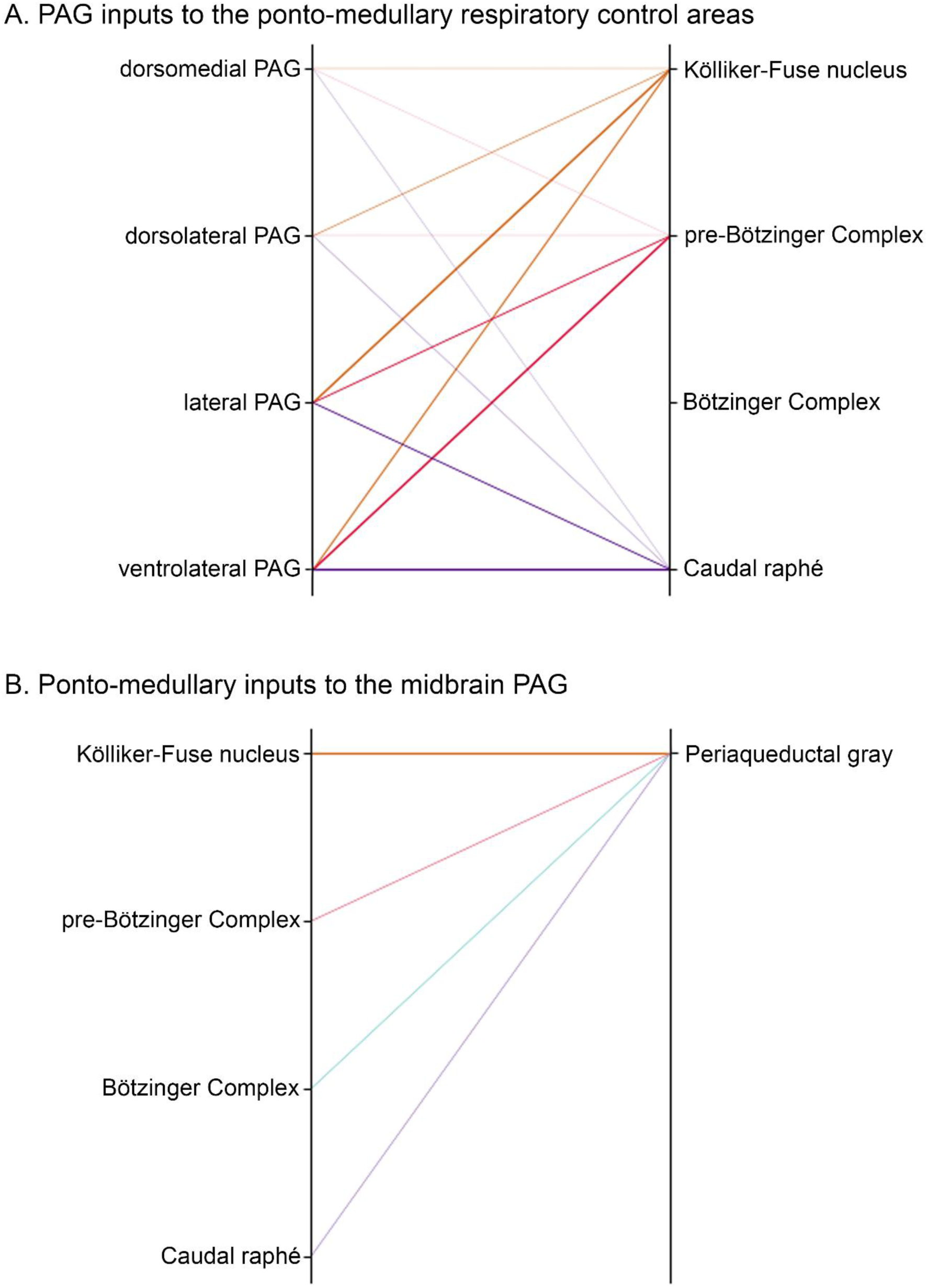
Summary of the relative strength of projections arising from the midbrain PAG columns (dmPAG, dlPAG, lPAG, and vlPAG; **A**), and the ponto-medullary regions (KFn, pre-BötC, BötC, and the caudal raphé; **B**) in a connectivity map. The weight of connecting lines are proportional to the normalized maximal number of CT-B-labeled neurons found in the specific source of descending projections. *Abbreviation*: PAG = periaqueductal gray.

## 3. Discussion

The midbrain PAG is thought to play a critical role in modulating breathing via inputs to the ponto-medullary respiratory network. However, PAG anatomical connectivity with core nuclei of the respiratory network remains vague. The present study identified neurons in the PAG that target either the pontine KFn, the caudal raphé, or the medullary pre-BötC, but not the BötC. It is likely that these descending PAG neurons are important for the volitional modulation or appropriation of respiratory motor activities. Several studies reported that stimulations of the PAG (lPAG and vlPAG, especially) evoke strong respiratory modulations and, in particular, trigger vocalizations (Magoun et al., 1937; Jürgens and Pratt, 1979, Larson and Kistler, 1984; Jürgens, 1994; Zhang et al., 1994; Davis and Zhang, 1996; Huang et a., 2000; Hayward et al., 2003; Zhang et al., 2007; Subramanian et al., 2008b; Jürgens, 2009; Zhang et al., 2009; Subramanian et al., 2013; Subramanian and Holstege, 2013; Farmer et al., 2014). Furthermore, changes in respiratory activity (i.e., an increase in the respiratory frequency) can also be correlated with defensive behaviors (e.g., fight and flight, freezing, aggressive or escape actions) that are elicited by PAG stimulation (Bandler, 1982; Bandler et al., 1985; Bandler and Depaulis, 1991; Carrive et al., 1997; Fanselow, 1991; Walker and Carrive, 2003; Hayward et al., 2003). The present study identified approximately equal numbers of PAG neurons projecting to the KFn, pre-BötC, and caudal raphé nuclei, which have different functions in respiratory pattern formation (*KFn*; see Dutschmann and Herbert, 2006; Smith et al., 2007; Mörschel and Dutschmann, 2009), rhythm generation (*pre-BötC*; see Smith et al., 1991; Wenninger et al., 2004; Feldman and Del Negro, 2006), and modulation of the excitability of respiratory network neurons (*caudal raphé*; see Lindsey et al., 1998; Richter et al., 2003, Hodges and Richerson; 2010). Taken together, our anatomical finding suggests that the PAG may engage a widely distributed network of brainstem nuclei to modulate respiratory motor pattern.

It has been proposed that the PAG might be the final common pathway mediating changes in the eupneic respiratory motor pattern in the context of various volitional and emotional behaviors (Subramanian and Holstege, 2014), including vocalization (Jürgens and Richter, 1986; Holstege, 1989; Jürgens 1994; Jürgens 2002; Düsterhöft et al., 2004; Subramanian and Holstege, 2009). However, the proposed descending pathway that connects the PAG output with laryngeal and spinal respiratory motor neurons was via solely on PAG projections to the nucleus retroambiguus (NRA) in the caudal medulla (Holstege et al., 1989; Holstege et al., 1997; Vanderhorst et al., 2000). In the PAG-NRA hypothesis, the NRA subsequently acts as an interface that distributes PAG signals to laryngeal motor neurons of the nucleus ambiguus (Vanderhorst et al., 2001) and various respiratory motor pools in the spinal cord (Vanderhorst and Holstege, 1995; Holstege et al., 1997), bypassing the respiratory rhythm and pattern generating network in the ponto-medullary brainstem, and is thought to be critical for PAG-mediated modulation of breathing, particularly during vocalization (Zhang et al., 1995; Subramanian and Holstege, 2009). However, recent work from our laboratory has shown that the eupneic respiratory motor pattern is generated by an anatomically widely-distributed brainstem network (Dhingra et al., 2019a,b; Dhingra et al., 2020). Consistent with these observations, the present study shows that the lPAG and vlPAG send multiple descending projections to these widely-distributed and functionally diverse brainstem respiratory control nuclei. For instance, in line with a previous study, the PAG receives afferents from the respiratory rhythm generating circuits of the pre-BötC (Yang and Feldman, 2018), and, therefore, may receive information regarding the respiratory rhythm. However, compared to pre-BötC, we observed that the KFn has significantly stronger afferent inputs to the PAG. Thus, the lPAG and vlPAG columns reciprocally connected with the KFn, which also receives substantial afferent input from the NRA (Jones et al., 2016).

In a recent study of the descending forebrain projections targeting the brainstem nuclei important for breathing control, we reported efferent cortical projections predominantly target the PAG and KFn (Trevizan-Baú et al., 2020). The reciprocal connectivity of the PAG and NRA with the KFn seems a coherent anatomical structure for the mediation of volitional respiratory modulation (e.g., vocalization and sniffing; see Trevizan-Baú et al., 2020), which requires control of expiratory airflow via glottal adduction during the post-inspiratory phase (Dutschmann et al., 2014). In contrast, changes in inspiratory frequency associated with emotion (Homma and Masuko, 2008) or behavior (Subramanian et al., 2008a, b; Subramanian and Holstege, 2014; Faul et al., 2019) might be mediated via forebrain to PAG projections (this study; Trevizan-Baú et al., 2020) targeting the pre-inspiratory rhythm generating circuit of the pre-BötC (Subramanian et al., 2013).

In conclusion, the midbrain PAG projects to the ponto-medullary respiratory network (i.e., pontine KFn, medullary pre-BötC, and the caudal raphé), and receives ascending projections from those respiratory control areas, especially from the KFn. The reciprocal PAG-KFn connectivity provides an anatomical framework for the regulation of upper-airway patency during the mediation of breathing modulation in the context of volitional and emotional behaviors.

## 4. Experimental procedures

### 4.1. Animals

Adult Sprague-Dawley rats of either sex (*n*=18, weight range: 280-350g) were used. Rats were housed under a 12:12 h light/dark cycle, with free access to lab chow (Ridley Corporation Limited, Australia) and water. All experimental procedures were approved by the Florey Institute of Neuroscience and Mental Health Animal Ethics Committee and performed in accordance with the National Health and Medical Research Council of Australia code of practice for the use of animals for scientific purposes.

### 4.2. Surgery

Details of the full surgical procedures are published in a recent study (Trevizan-Baú et al., 2020) In brief, after craniotomy, a 1 µl Hamilton syringe (25s-gauge needle) filled with 150nL of 1% CT-B (1 mg/mL; Invitrogen, OR, USA) was used to pressure inject the retrograde tracer (150nL; 20 nL/min) unilaterally (left side of the brain) in the KFn, BötC, pre-BötC, caudal raphé nuclei, or PAG. We used the following coordinates (Paxinos and Watson, 2007): KFn: -9 mm relative to bregma (A/P), 2.6 mm lateral to the midline (M/L), and 6.5 mm ventral to the dorsal surface of the brain (D/V); BötC: -9 mm A/P, 2 mm M/L, and 9.5 mm D/V; pre-BötC: -9.5 mm A/P, 2 mm M/L, and 9.5 mm D/V; caudal raphé nuclei: -8.7 mm A/P, 0 mm M/L, and 9.4 mm D/V; and PAG: -8 mm A/P, 0.8 mm M/L, and 4.8 mm D/V. Injections in the medullary regions (i.e., BötC, pre-BötC, and caudal raphé), required an adaptation and the syringe was angled at 22° backwards from vertical to avoid bleeding from superficial cerebral blood sinuses. After injections were completed, the syringe remained in the brain tissue for 10 min and was withdraw at 1 mm/min to minimize non-specific spread of the tracer along the injection tract (Finkelstein et al, 2000).

### 4.3. Tissue preparation and immunohistochemistry

The full experimental protocols for the immunohistochemistry are published in Trevizan-Baú et al. (2020). In brief, 12-14 days post-injections, rats were transcardially perfused with paraformaldehyde (PFA, Sigma-Aldrich) containing 0.2% picric acid (Sigma) diluted in 0.16 M phosphate buffer (Merck KGaA, Darmstadt, Germany; pH 7.2, initially at 37 °C and subsequently at 4 °C). After postfixation with PFA/picric acid solution for 90 min at 4°C, the brains were immersed for 72 h in 0.1 M phosphate buffer (pH 7.4) containing 15% sucrose, 0.01% sodium aside (Sigma) and 0.02% bacitracin (Sigma) for cryoprotection. Before brain were sectioned with a cryostat (Leica CM1850, Leica Microsystems; coronal sections of 40 µm thickness), a small superficial incision was made from the midbrain to the brainstem to distinguish the ipsilateral and contralateral sides of the brain relative to the injection site. Free-floating sections were washed in 0.01 M PBS, followed by incubation in hydrogen peroxide for 20 min and incubated for 24 h at room temperature (RT) with a goat anti-CT-B antibody (1:10,000 List Biological Laboratories, CA, USA). Then, sections were washed in 0.01 M PBS, blocked with 5% normal donkey serum (NDS) in 0.01M PBS for 1 h at RT, incubated in the corresponding secondary antibody (anti-sheep biotin, Jackson ImmunoResearch Laboratories, West Grove, PA; 1:500 in 0.01 M PBS) for 1h at RT, and subsequently incubated in an ABC kit (Vectastain® Elite ABC-HRP Kit) for 1h at RT. Finally, sections were incubated in diaminobenzidine (DAB) substrate (1:10, Roche Diagnostics Mannheim, Germany) for 4 min, followed by 1% H2O2 incubation for 4 min. After several washing steps, the sections were then mounted on slides and cover-slipped.

### 4.4. Data analysis and image processing

To identify and document the location of the midbrain and brainstem injection sites, we used a brightfield microscope (Leica Biosystems). Injection sites were photographed and represented on a schematic drawing (Fig. 1A; Fig. 3A). Additional information regarding the distribution and location of the injection sites are detailed in our previous study (Trevizan-Baú et al., 2020).

For analysis, we quantified the number of CT-B-labeled neurons in the brain region of interest. CT-B-labeled neurons were recognized by the presence of black punctate granules in the neuronal cell bodies (somata) and dendrites, and the absence of stained cell nuclei (for details, see Trevizan-Baú et al., 2020). Cell counts were analyzed with GraphPad Prism (version 7.02; GraphPad Software; San Diego, CA). Data were analyzed by One-way ANOVA followed by Tukey’s multiple comparison test to compare the number of CT-B-labeled neurons between the target regions. To determine statistical significance between the numbers of CT-B-labeled neurons in the ipsilateral and contralateral hemisphere, data were analyzed by paired t-test (parametric). All values are presented as mean ± standard error of the mean (SEM), and *p*-values less than 0.05 were considered statistically significant.

A digital camera (Leica DFC7000T) mounted to the microscope (Leica DM6B LED) was used to take representative images. To best represent sections viewed under the microscope, we used the Adobe Photoshop CC19 software (Adobe Systems Inc., San Jose, CA) to optimize brightness and contrast of the digital images, as well as to assemble representative images.

## Acknowledgements

We acknowledge that this work was conducted on the traditional land of the Wurundjeri people of the Kulin nation. We pay our respect to their elders past, present and emerging. This work was supported by research grants from the National Health and Medical Research Council of Australia (APP1165529 to MD & DS, and APP1078943 to SM); and Australian Research Council (Discovery project DP170104861); PT-B is funded by Melbourne Research Scholarship (University of Melbourne; 181858).

## Author contributions

PT-B, DS and MD conceived and designed the experiments. P-TB and MD analyzed the data and prepared figures and tables. PT-B conducted all experiments. All authors contributed to interpretation of the data, reviewed drafts of the manuscript, and approved the final draft.

